# Tracing diagnosis trajectories over millions of inpatients reveal an unexpected association between schizophrenia and rhabdomyolysis

**DOI:** 10.1101/473082

**Authors:** Hyojung Paik, Matthew J. Kan, Nadav Rappoport, Dexter Hadley, Marina Sirota, Bin Chen, Udi Manber, Seong Beom Cho, Atul J. Butte

**Affiliations:** Bakar Computational Health Sciences Institute, University of California, San Francisco, 550 16th Street, San Francisco, CA 9414, USA; Department of Pediatrics, University of California, San Francisco, 550 16th Street, San Francisco, CA 94143, USA; Korea Institute of Science and Technology Information, Center for Applied Scientific Computing, Division of Supercomputing, Daejeon 34141, South Korea; Department of Medicine, University of California, San Francisco, 505 Parnassus Avenue, San Francisco, CA 94143, USA; National Institute of Health, Division of Bio-Medical Informatics, Center for Genome Science, OHTAC, 187 Osongsaengmyeong2(i)-ro, Gangoe-myeon, Cheongwon-gun, ChoongchungBuk-do, South Korea

## Abstract

While it has been technically feasible to create longitudinal representations of individual health at a nationwide scale, the use of these techniques to identify novel disease associations for the risk stratification of patients has had limited success. Here, we created a large-scale US longitudinal disease network of traced readmission patterns (i.e., disease trajectories), merging data from over 10.4 million inpatients from 350 California hospitals through the Healthcare Cost and Utilization Project between 1980 and 2010. We were able to create longitudinal representations of disease progression mapping over 300 common diseases, including the well-known complication of heart failure after acute myocardial infarction. Surprisingly, out of these generated disease trajectories, we discovered an unknown association between schizophrenia, a chronic mental disorder, and rhabdomyolysis, a rare disease of muscle breakdown. It was found that 92 of 3674 patients (2.5%) with schizophrenia were readmitted for rhabdomyolysis (relative risk, 2.21 [1.80–2.71, confidence interval = 0.95] *P*-value 9.54E-15), which has a general population incidence of 1 in 10,000. We validated this association using independent electronic health records from over 830,000 patients treated over seven years at the University of California, San Francisco (UCSF) medical center. A case review of 29 patients at UCSF who were treated for schizophrenia and who went on to develop rhabdomyolysis demonstrated that the majority of cases (62%) are idiopathic, which suggests a biological connection between these two diseases. Together, these findings demonstrate the power of using public disease registries in combination with electronic medical records to discover novel disease associations.

**One Sentence Summary:** Based on the longitudinal health records from millions of California inpatient discharges, we created a temporal network that enabled us to understand statewide patterns of hospital readmissions, which led to the novel finding that hospitalization for schizophrenia is significantly associated with rehospitalization for rhabdomyolysis.

## Introduction

One of the most basic aspects of clinical care is understanding disease and mortality risk factors, particularly for rare but preventable diseases or outcomes (Ashley et al., 2010; Camilo and Goldstein, 2004; Finkelstein et al., 2009). Although mapping disease relationships has a long history, the recent advent of digitalized health records and disease registries has led to an enhanced ability to organize and analyze healthcare data. The availability of these data allows the recapitulation of known temporal disease correlations using nonlongitudinal data (Hidalgo et al., 2009), exploration of unordered disease pairs (Blair et al., 2013; Park et al., 2009) and enabling network analyses of disease relationships at a national scale (Jensen et al., 2014). However, while big-data analytics have aided the ability to visualize, search, and organize health data (Murdoch et al., 2013; Yeo et al., 2013), there has been limited success in identifying and validating novel temporal relationships between diseases that would meaningfully change clinical care.

Here, we showcase a novel temporal disease association discovered in the course of a large-scale analysis of California inpatient hospitalizations using data from the Healthcare Cost and Utilization Project (HCUP) (Steiner et al.). By using the California State Inpatient Database (CA SID) from HCUP, we analyzed longitudinal inpatient discharge data collected between 1980 and 2010 from 350 California hospitals, representing 10.4 million ethnically diverse individuals. Based on inpatient diagnoses, we identified temporal correlations between disease pairs (Jensen et al., 2014) and concatenated these correlations to create a network-based representation of disease trajectories from initial hospitalization until death in a hospital. By using a combination of computational analysis and healthcare expert curation, we discovered an unexpected relationship between inpatient admissions for schizophrenia, a psychiatric disorder, and readmission for rhabdomyolysis, a rare disease of muscle breakdown. We validated this finding by performing a case review of patients treated for schizophrenia and rhabdomyolysis at the Medical Center of the University of California, San Francisco (UCSF), thereby demonstrating the reproducibility of the identified association between schizophrenia and rhabdomyolysis. Together, these findings demonstrate the power of using multi-level medical records including public disease registries in combination with detailed electronic medical records to discover novel disease associations, explore their etiologies, and inform clinical practice.

## Results

### Data set and overview of analytics

As a main study set, we used the five editions of the annually released HCUP datasets from the CA SID, which contains inpatient billing records organized by the Agency for Healthcare Research and Quality (AHRQ) (https://www.hcup-us.ahrq.gov), including *International Classification of Diseases, Ninth Revision*, Clinical Modification (ICD-9-CM) diagnosis codes and death outcomes in hospitals, collected from over 350 community hospitals in California. The CA SID uses unique patient identifiers for each edition; thus, discharge data within the years is kept together for each patient within the edition, such as between 1980 and 2010 in the 2010 edition, but cannot be linked across previous editions. Therefore, while each edition of CA SID covers the longitudinal history of inpatient records for each patient across over 20 years (Figure 1A), the patient record cannot be mapped between different editions of CA SID. To prevent recounting the same patients, we started with the 2010 data edition and then added record data from previous editions only for patients that were deceased during that year (and thus could not have been admitted again in the 2010 edition) (See Methods and **Figure 1A**). The patients may be treated for multiple illnesses during a hospitalization, both acute and chronic, so the first assigned diagnosis was used as a principal diagnosis for each admission. Given that we were interested in tracing the relationships of diseases, we filtered out ICD codes for nondisease conditions, which we defined as diagnosis chapters from ICD-9-CM, referring to injury, pregnancy, external causes, and healthcare-related administration codes. The merged data set covered 2,272,018 hospitalizations for 1,488,551 individuals with 290,253 death outcomes. The mean age of patients in the merged CA SID was 63.77 ± 19.58 years (**Supplementary table S1**).

**Figure 1:**
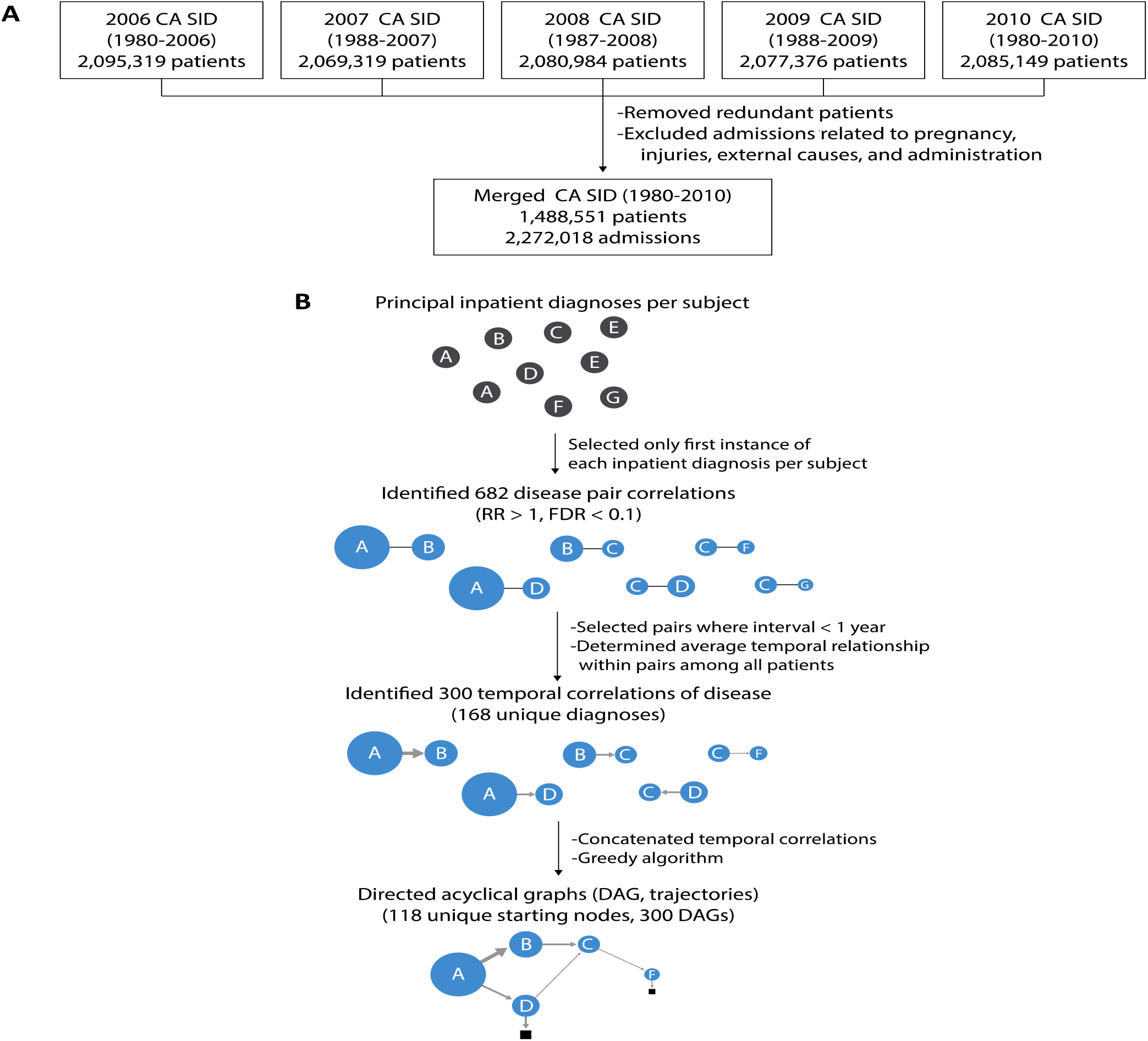
Overview of a main study data and analytics. (A) Overview of the data set build and analytics. We prepared the study data set by combining five editions of the California State Inpatient Database (CA SID) sets, which were released annually (2006–2010). In summary, we removed admissions for which primary diagnoses were not related to disease (pregnancy, injury, external causes, administration). The merged CA SID covers longitudinal records (1980–2010) for >1.4 million patients without data redundancy. (B) Construction of DAGs. The principal inpatient diagnoses for patients were selected and 682 significant disease association pairs (RA > 1, FDR < 0.1) were identified. The average temporal correlation was determined within pairs and then concatenated by using a greedy algorithm to create DAGs (i.e. trajectories) with 117 starting nodes.

As we were interested in temporal relationships between disease diagnoses within individuals, we simplified the timeline of each patient to the first instance of each primary diagnosis, such that readmissions for the same disease were removed. All diagnosis codes were reported using the ICD-9-CM and rounded to the three-digit code level (O’Malley et al., 2005) to minimize overlap and subclassification of diagnoses (see Methods). Based on the intervals between the first and last admission dates for each individual, most of the merged CA SID data included diagnostic timelines for each patient of over three years (the median interval between the first and last admission dates per patient was 40.58 ± 2.5 months). The longest duration between the first and most recent admission dates in a patient was 26 years. We then identified significant disease associations by establishing the relative association (RA) of diagnoses co-occurring within one year in a patient (Hidalgo et al., 2009; Park et al., 2009) using a binomial test, and compensating for multiple hypothesis testing using a false discovery rate (FDR) < 0.1 (see Methods) (Jensen et al., 2014). In short, RA is a ratio of the actual probability of codiagnoses of two diseases over the random expectation of co-occurrences. When RA is greater than 1, it indicates the observed disease co-occurrence is higher than expected by chance (the details are shown in the Methods). We chose an interval of one year because the majority of new admissions of patients occurred within the same year, and admissions separated by very long-term intervals were less likely to be related.

Of the 691 three-digit level ICD-9-CM diagnosis codes used in the merged data set, 168 disease diagnoses were associated with at least one of the other 167 diseases, with a total of 300 temporally aligned disease pairs at FDR < 0.1. We determined the average time direction within each pair to determine the temporal relationship between diagnoses. Given that our data (CA SID) represent a statewide inpatient data set, the initial date of disease diagnosis is based on the date of admission for a disease. By using a greedy search algorithm (Jensen et al., 2014) (i.e., traced temporal relationships between diagnoses with the largest number of patients), we then concatenated these 300 temporal parings to create larger directed acyclic graphs (DAGs). By iterating over each starting disease built DAGs, we retrospectively mapped 300 common trajectories from initial hospitalization through intermediate hospitalizations until death, if applicable (**Figure 1B**). The 300 presented DAGs have 118 different diseases as nodes for the first diagnosis. The associated admissions cover 311,309 patients, who had 175,556 recorded readmissions for distinct diseases that are statistically correlated and that are presented as edges between disease diagnosis nodes. Of the traced patients, 34% (59,794) reached fatal outcomes as hospital inpatients.

### DAGs depict temporal disease associations and mortality

In total, we identified 300 major trajectories (i.e. DAGs) consisting of nodes for disease diagnoses, and edges for subsequent readmission for statistically associated disease diagnoses. Of the 300 trajectories, the longest trajectory had four readmission steps for correlated diseases from the initial presentation of disease diagnosis. In 257 trajectories, the latest steps of readmissions are associated with fatal outcomes, i.e., deaths in clinics. While the dataset is too comprehensive to review all of our findings in detail (see **Supplementary Movie 1** for overview of overall findings), we highlight a few selected trajectories (i.e. DAGs). We focused on three DAGs that represent the disease trajectories leading to the most hospital deaths of California within the network (**Figure 2**). Among patients in their fifties, most hospital deaths (2,237 deaths) occurred within the DAG associated with a “chronic liver disease and cirrhosis,” which was the initial presentation of disease in the DAG for 5416 patients. A significant portion (up to 35%) experienced infection complications (liver abscess and sequelae of chronic liver disease) and then sepsis, with most deaths associated with these ICD diagnoses (**Figure 2A**). We also recapitulated a well-known phenomenon that acute myocardial infarction commonly leads to heart failure and that these steps of diseases have high mortality (19%; 2,245 deaths among 11,624 of the traced acute myocardial infarction patients) (**Figure 2B**). Thus, identified diagnosis patterns within DAGs appeared to be reliable and represent currently understood phenomena. We also present the DAG for pneumonia, which disproportionately affects the elderly (mean age in the 70s). This DAG demonstrated a network with both high morbidity and mortality. Associated subsequent diagnoses included heart failure and subsequent lung disease, aspiration pneumonia (pneumonitis due to solids and liquids), and sepsis (**Figure 2C**). Indeed, sepsis was a common dead-end node in many DAGs, which reinforces previous studies demonstrating the high prevalence of sepsis in the USA (Martin, 2012). Together, these findings allow common disease paths to be traced and enables diseases of high mortality in clinics to be identified in a systematic fashion. In addition to recapitulating known disease trajectories, greater detail of 300 trajectories (i.e. DAGs) are presented in **Supplementary Data S1**.

**Figure 2:**
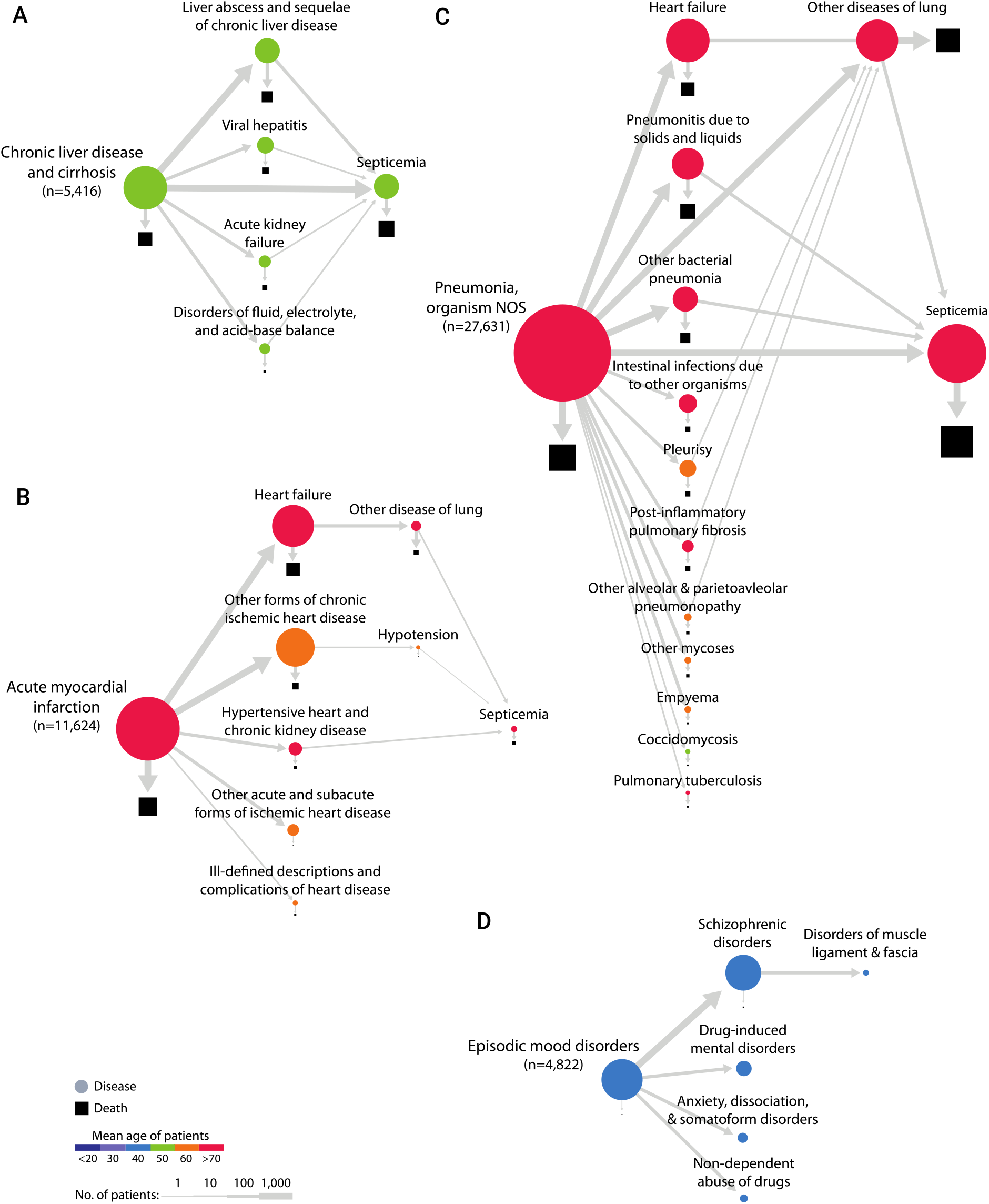
Selected disease trajectories (DAGs). Selected DAGs (117 in total) showing trajectories for one-year intervals between primary diagnosis codes for California inpatient admissions. Areas of shapes are directly proportional to the number of patients, where circles represent primary diagnoses and squares represent deaths. Nodes are colored by mean age of patients. The thickness of edges is determined by the number of patients traveling along the edge (see legend). (A) DAG of “chronic liver disease and cirrhosis.” Prominent nodes include “liver abscess and sequelae of chronic liver disease,” “septicemia,” and “death.” (B) DAG of “acute myocardial infarction.” The majority of readmissions were for “heart failure” and “other forms of chronic ischemic heart disease,” and “acute myocardial infarction” and “heart failure” were the nodes most associated with death. (C) DAG of “pneumonia, organism NOS.” Common readmissions were for “septicemia,” “heart failure,” “other disease of lung,” “pneumonitis due to solids and liquids,” and “other bacterial pneumonia.” These diagnoses were all strongly associated with death. (D) DAG of “episodic mood disorders,” which includes diagnosis codes for uncharacterized mental disorders. The majority of hospitalizations were readmitted for diagnoses of “schizophrenia,” and a significant proportion of these individuals were later hospitalized for “disorders of muscle ligament and fascia;” 93% of the admissions for “disorders of muscle ligament and fascia” were more specifically coded for rhabdomyolysis (ICD-9-CM code = 728.88).

### Schizophrenia is associated with subsequent hospitalization for rhabdomyolysis

We also found a surprising relationship between schizophrenia and muscle disorders (**Figure 2D**). In this DAG, 4,822 patients were first diagnosed with episodic mood disorders, which are often used in initial diagnoses for psychiatric diseases while medical providers collect collateral data to make a more specific clinical diagnosis. As expected, 76% (3,674) were subsequently readmitted with schizophrenia (schizophrenic disorders). While diagnoses for psychotic disorders exist on a continuum of clinical symptoms, the fifth edition of the *Diagnostic and Statistical Manual of Mental Disorders* (DSM-V) requires either two outpatient evaluations or one inpatient hospitalization to make a diagnosis of schizophrenia. Therefore, we felt confident in this categorization. Interestingly, 98 of the 3,674 patients (2.6%) with schizophrenia were readmitted within one year with muscle disorders. Of these 98 patients with disorders of muscle, ligament, and fascia, 92 (2.5% of 3,674 schizophrenia patients) were more specifically coded as having rhabdomyolysis, which is a very rare disease of muscle breakdown that can lead to kidney failure and even death (relative risk [RR] = 2.21 [1.80–2.71, confidence interval (CI) = 0.95] *P*-value of RR 9.54E-15, RA = 1.5, mean interval 114.9 ± 84.3 days, FDR of RA and temporal order = 8.76E-02) (**Figure 3A**). Although it has been historically very difficult to study the epidemiology of rhabdomyolysis, an incidence of lower than 0.0001 (26,000/year among the 325 million US population, 8E-05 of incidence per year) has been estimated (American Academy of Family Physicians et al., 1970). Thus, a rate of 2.5% in a specific patient population (schizophrenia) was peculiarly high (313-fold higher than a population-wide incidence of rhabdomyolysis; 0.026 is 325 times greater than 8E-05) (*P*-value of enrichment using hypergeometric test 7.20E-03). There has been one case report describing second generation antipsychotics, such as aripiprazole (Chang and Wu, 2011) (a schizophrenia drug for adolescence patients (Greenaway and Elbe, 2009)), causing rhabdomyolysis in a schizophrenia patient. Regarding the rare and immediate adverse effect of aripiprazole, the association in middle-aged adults (mean age, 40.39 years) is not a widely known disease association (mean of intervals between schizophrenia and rhabdomyolysis = 114.9 ± 84.3 days). Thus, we sought to verify and further characterize this novel relationship.

**Figure 3:**
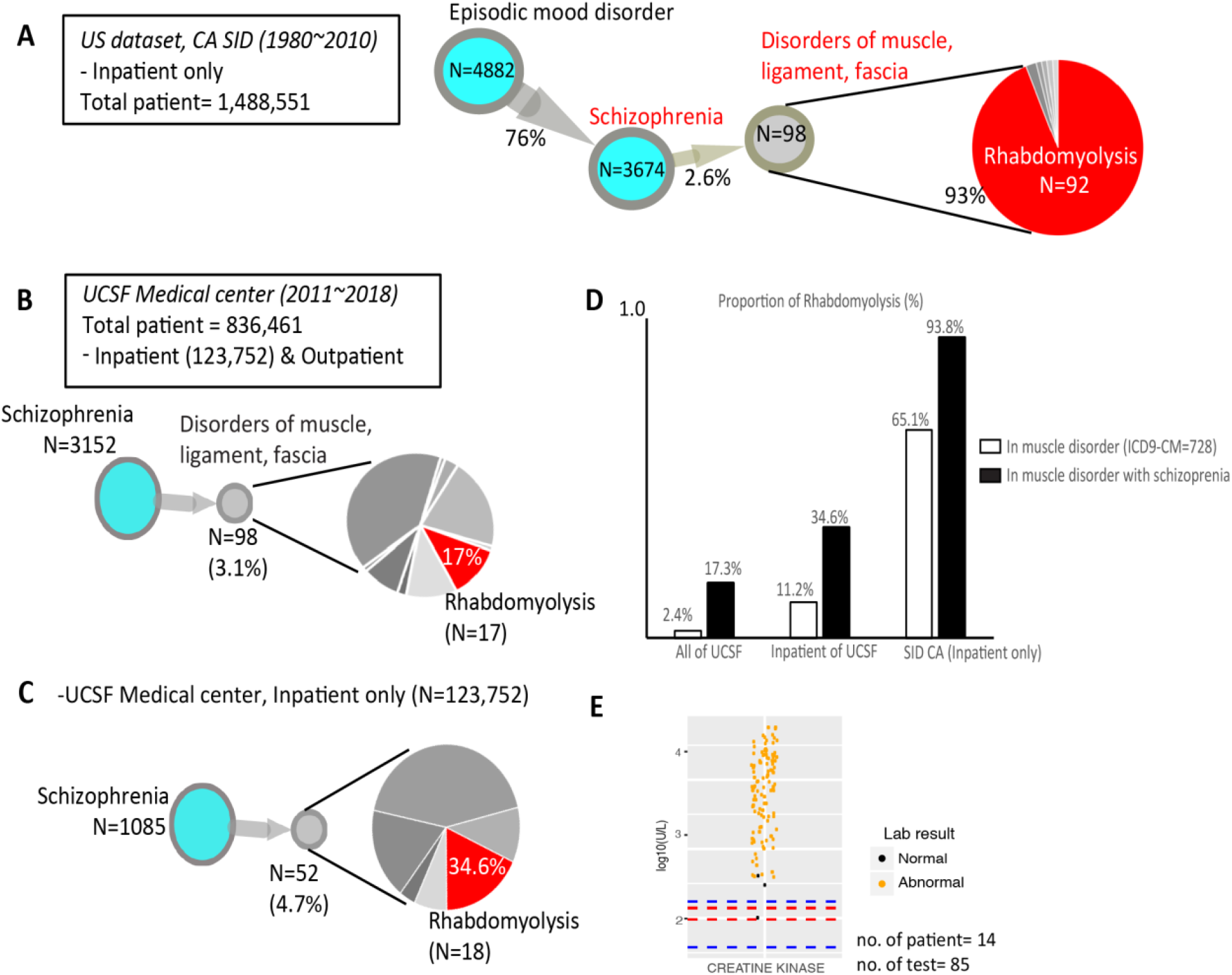
The identified novel association between schizophrenia and rhabdomyolysis. Our scaled analysis of digitalized medical records from millions of patients reveals a novel association between schizophrenia and rhabdomyolysis. (A) DAG of schizophrenia and disorders of muscle, ligament, and fascia enriched with rhabdomyolysis based on the data of CA SID. (B) DAG of schizophrenia and disorders of muscle, ligament, and fascia enriched with rhabdomyolysis based on the EHRs of the UCSF Medical Center. (C) The identical DAG of (B) using the subset of the EHRs of the UCSF Medical Center consisting of inpatient records. (D) The proportion of the diagnoses of rhabdomyolysis in schizophrenia across different data sets. Black bars indicate the fraction of rhabdomyolysis among the muscle disease (ICD-9-CM code = 728) in schizophrenia. White bars represent the rate of rhabdomyolysis in muscle disease-diagnosed patients. (E) Lab test evidence for the diagnoses of rhabdomyolysis in (B). The levels of creatine kinase in patients who were diagnosed with rhabdomyolysis after schizophrenia are extremely high. The blue and red lines are the reference range for normal levels of creatine kinase (38–174 units/L [reference for male, blue line]; 96–140 units/L [reference for female, red line]).

### Multi-level validation of patients with schizophrenia diagnosed with rhabdomyolysis

To better understand the potential etiology of rhabdomyolysis in schizophrenia patients, we searched for this relationship using the electronic health records (EHRs) of the UCSF medical center, which is one of the contributors to the CA SID. At the time of writing, the UCSF EHR data consists of inpatient and outpatient encounters from 836,461 patients between 2011–2018 (inpatients = 123,752 patients), whereas merged CA SID consisting of five editions of CA SIDs covers inpatients from 1980 to 2010. Thus, the data of UCSF and CA SID rarely overlap. We began by using UCSF’s deidentified Clinical Data Warehouse of EHRs, which contains structured data including diagnoses, lab values, vitals, procedures, and medication administrations, without any direct identifiers.

Similar to CA SID data, we found that the proportion of UCSF patients with schizophrenia who were treated for “disorders of muscle ligaments and fascia” to be statistically higher than expected (RA = 1.1, mean interval = 296.7 ± 311.1 days, FDR < 0.1). When we tracked these data in a time-ordered manner, 98 of 3,152 schizophrenia patients (3.1%) were hospitalized or visited for muscle disorder after the schizophrenia, and 17 of those 98 (17.3%) were enriched with rhabdomyolysis (*P*-value of hypergeometric test 9.9E-22) (**Figure 3B**). In UCSF, the number of rhabdomyolysis patients was 541 among 836,461 patients (0.06%). Thus, in the UCSF Medical Center, the diagnosis of rhabdomyolysis is peculiarly high among schizophrenia-diagnosed patients (0.53%, 17 of 3152, RR = 18.3 [13.4–25.0, CI = 0.95] *P*-value of RR = 2.2E-16). A similar pattern was captured among 1,085 schizophrenia inpatients of the UCSF Medical Center (*P*-value of enrichment using hypergeometric test 1.51E-11) (**Figure 3C**) (1.65%, 18 of 1085, RR = 9.61 [6.8–13.4, CI = 0.95] *P*-value of RR=1.1E-15).

In addition, the enrichment patterns of rhabdomyolysis diagnoses among the muscle disorders in schizophrenia are repeated across the data of UCSF EHRs and CA SID (**Figure 3D**). Among 22,374 muscle disease patients in our UCSF data set (ICD-9-CM code ‘728’ for disorders of muscle, ligament and fascia), only 541 (2.4%; shown as a white bar in Figure 3D) were rhabdomyolysis (ICD-9-CM code ‘728.88’). However, the diagnosis fraction of rhabdomyolysis was sevenfold higher among the overall muscle diseases in schizophrenia patients in UCSF (17.3%, 17 of 98 muscle disease patients with schizophrenia, shown as a black bar). The black bars in Figure 3D represent the proportions of patients that were diagnosed as rhabdomyolysis among patients both diagnosed as muscle disorder and schizophrenia. These proportions were larger than the fraction of rhabdomyolysis in overall muscle disorder in UCSF EHRs and CA SID, which are indicated by the white bars.

We also examined whether rhabdomyolysis occurred following schizophrenia due to the adverse effect of aripiprazole, a second-generation antipsychotic (Chang and Wu, 2011); however, none of the schizophrenia inpatients of the UCSF Medical center were administrated aripiprazole. Other antipsychotic drugs, including haloperidol and clozapine, have less clear association with rhabdomyolysis (*P*-value of odds ratio >0.05); the association between schizophrenia and rhabdomyolysis in UCSF is thus independent of adverse drug effects.

Diagnoses of rhabdomyolysis for billing purposes including healthcare cost were also validated based on clinical evidence (lab values). The levels of creatine kinase for the diagnostic confirmation of rhabdomyolysis greatly exceeded the normal reference (**Figure 3E**) (normal reference of creatine kinase = 38–174 units/L (reference for male, blue line in **Figure 3E**); 96–140 units/L (reference for female, red line in **Figure 3E**)) in those of schizophrenia–rhabdomyolysis patients (mean = 9137.74 ± 14808 units/L), which allowed a high confidence in the correct diagnoses of rhabdomyolysis occurrence in patients with schizophrenia (14 of 17 patients in Figure 3B).

For the protection of privacy, the deidentified EHR of UCSF consists of structured types of data, such as diagnosis code. For detailed case reviews, after obtaining approval from the Institutional Review Board of UCSF, we recollected the data from 29 cases in which rhabdomyolysis was diagnosed after schizophrenia, from all possible clinical data warehouses in UCSF (approval number 17-22258). A case review of physicians’ notes for patients with schizophrenia treated for rhabdomyolysis at UCSF between 2011 and 2018 (*n* = 29) revealed that 37% of these cases (*n* = 11) involved illicit drug ingestions, and 13% (*n* = 4) were also from fractures/falls. Other muscle symptoms in schizophrenia that can be relevant to rhabdomyolysis, such as neuroleptic malignant syndrome (NMS), catatonic seizure and spasm, were not detected from the healthcare records including physicians’ notes on these subsets of schizophrenia patients who were diagnosed with rhabdomyolysis at UCSF (Melli et al., 2005). However, over half were of idiopathic origins (*n* = 18, 62%), and had no clinically attributable cause. This could suggest that there could be a shared risk factor or potential biological predisposition for rhabdomyolysis with underlying schizophrenia. All of the case review processes were conducted by professional medical providers, including psychiatric physicians.

## Discussion

From the population-scale analysis of digital health records, the unexpected propensity of rhabdomyolysis to occur in patients with schizophrenia (0.5–2.5%) was consistently detected compared with random and sporadic chances across the two data sets composed of tens of millions of individuals (CA SID and UCSF EHRs, RRs 2.21–18.35). Known possible risk factors for rhabdomyolysis, such as adverse effects of medication and NMS, were not reported in the majority of rhabdomyolysis cases following schizophrenia. Thus, our national scaled analysis of digitalized medical records from millions of patients reveals a novel association between schizophrenia and rhabdomyolysis via shared idiopathic mechanisms.

This study included an extensive temporal analysis of disease occurrences across the discharge records of 10.4 million patients in Californian hospitals. In total, we identified 300 serial readmissions for diverse disease diagnoses beginning with 118 disease nodes in 311,309 patients. Throughout 175,556 succeeding admissions with temporally correlated diagnosis from previous admissions and a relevant diagnosis, our approach presents time-aligned patterns of readmission and consistently modeled known clinical associations, such as heart failures following acute myocardial infarction. Interestingly, from the modeled disease trajectories, we repeatedly identified a novel association of diseases including rhabdomyolysis following schizophrenia from the independent data set. By combining multilevel validation and case reviews of these patients with professional medical providers, an unexpected risk of rhabdomyolysis in schizophrenia was revealed. This finding constitutes direct evidence for the value of digital health records for patient care in influencing clinical practice in a systematic fashion, such as identifying the risk of further disease.

It is known that patients with schizophrenia and other serious psychoses are underdiagnosed and undertreated for comorbid illnesses, such as hypertension and diabetes (Goldman, 1999). Thus, given the absence of the observational data recapitulation and time-ordered presentation, the association with rhabdomyolysis would have been overlooked. Potential etiologies include drug associations because patients with schizophrenia are often treated with antipsychotic medications, and there is a known relationship between second-generation antipsychotics causing NMS, a disease of catatonia and muscle spasm that can lead to muscle breakdown. There have also been rare case reports of second-generation antipsychotic use in patients with schizophrenia causing rhabdomyolysis in the absence of NMS. Patients with schizophrenia are more vulnerable to socioeconomic stressors, such as illicit substance use and homelessness, which are all independent risk factors for rhabdomyolysis. While these explanations are still more likely, there could also be biological explanations. There are known brain disorders including schizophrenia that share genetic etiologies with diverse diseases (Anttila et al., 2018). Over 108 genetic loci are associated with schizophrenia (Consortium et al., 2014). It is possible that an underlying genetic predisposition can affect both neurons and muscle cells. For example, AHI1 is associated with both schizophrenia and metabolic abnormalities of muscle (Prasad et al., 2017; Prior et al., 2010) and perhaps could contribute to these two disease predispositions.

Several limitations of the study should be noted. Other diverse and overlooked clinical and patient features are invisible in our data, such as neglected symptoms including mild muscle spasm, and unreported behaviors relevant to rhabdomyolysis after the discharge of patients (Life et al., 1997). In further studies, finer acquisition of EHRs for the computational approaches, such as natural language processing of the chart records, will help in the care of schizophrenia patients and facilitate risk assessment for rhabdomyolysis. However, based on our findings, the awareness of rare but critical risk (i.e., rhabdomyolysis) of schizophrenia patients may be helpful for providing adequate clinical practice. Further studies on the shared mechanistic or biological etiologies between schizophrenia and rhabdomyolysis should be conducted.

Two key messages can be deduced from this study. First, the analysis of health records can aid in advancing clinical knowledge for patient care. While most physicians are eager to comply with the latest evidence guiding clinical practice, scientific inquiries have been slow to uptake the rapidly expanding amount of data that are available from clinics. By using advanced data-mining analytics, we demonstrate how large-scale digital healthcare data can be used to facilitate the generation of medical knowledge. Second, the validation of the novel association between schizophrenia and rhabdomyolysis from the large-scale analytics depends mainly on the multilevel evidence of health records including finer validation of EHRs using lab test results and chart review with clinicians. Thus, for the specific contribution of big-data analytics and to effectively draw beneficial inference from clinical practice, communication with data scientists and physicians who have insight from routine clinical practice is indispensable. Although disease registries including CA SID are useful resources for observational studies, the use of a diverse range of evidence from clinical records, such as EHRs, are essential for novel findings. In this report, we have focused on a single example of our large-scale analysis (rhabdomyolysis after schizophrenia). To disseminate our findings, all of our results are available on the website, where the mapped associations of diseases can be explored via dynamic visualization applications such as Google map. Supplementary Movie S1 and Supplementary Data S1 present the prevision of the website (http://52.89.56.137:3000/#/intro, optimized for Chrome).

By tracking millions of healthcare records, this study has revealed a novel and consistent propensity for schizophrenia patients to develop rhabdomyolysis. Digital health data are a promising resource that can be used to create observational evidence for clinical questions, such as readmission for relevant diseases, and to identify novel risk of disease patients, which can influence clinical practice.

## Methods

### California administrative healthcare records dataset

We used the HCUP CA SID from the AHRQ. This database contains deidentified admission and discharge billing information from over 350 community hospitals in California, including nonfederal, general, specialty, and academic medical centers. The CA SID excludes noncommunity hospitals, such as federal hospitals (e.g., Veterans Affairs), long-term care hospitals, and clinical units within institutions (e.g., prisons).

We utilized five of the most recent annual builds of CA SID, which included 364 hospitals in edition 2006 (longitudinal inpatient records between 1980–2006), 360 hospitals in edition 2007 (records between 1988–2007), 361 hospitals in edition 2008 (records between 1987–2008), 354 hospitals in edition 2009 (records between 1988–2009), and 354 hospitals in edition 2010 (records between 1980–2010). We merged these five CA SID editions, each of which consists of over 20 years of longitudinal records for about 2 million patients. Thus, the merged CA SID covers inpatient records between 1980 and 2010. While each edition of CA SID used unique identifiers for every individual, allowing us to trace patients temporally across hospitals in a longitudinal manner, these identifiers were not consistent across versions, preventing meta-mapping of the patient identifiers between editions. To prevent data redundancy in the merged CA SID data set, we used records of only deceased individuals and their hospitalization records in the 2006–2009 editions, and merged these with the most recent 2010 edition.

All diagnosis codes were reported using the ICD-9-CM and rounded to the three-digit code level (O’Malley et al., 2005) to minimize overlap and subclassification of diagnoses. For example, the three-digit ICD-9-CM code for vascular dementia (F01) involves subtypes consisting of dementia with or without behavioral disturbances (F01.50 or F01.51); therefore, we used three-digit ICD-9-CM codes to avoid diagnosis subclassifications for vascular dementia.

### Temporal disease diagnoses correlations

We used the first charted diagnosis as a primary diagnosis code to determine the main reason for each patient hospitalization. To determine the temporal association of admission diagnoses correlations, we first calculated the RA measurement of all disease pairs (*Disease i – Disease j*) that occurred within one year for each patient (Park et al., 2009). Here, we used diagnoses for the patients based on the assigned three-digit ICD-9-CM code. We selected pairs where RA was greater than one, which indicated that the co-occurrence of the two diseases was higher than expected by chance. Then, by modifying a previous method, we determined the likelihood within a pair of diseases for one disease to occur before the other (*δ*_*i→j*_ for *disease i → disease j*) by using the dates of admissions associated with the two diseases in each patient (Hidalgo et al., 2009). To calculate *δ_i→j_*, we compared diagnoses dated for each patient. We defined 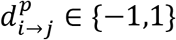 as 1 if disease *i* was diagnosed before disease *j* in patient *p* and −1 otherwise. Multiple rediagnoses or rehospitalizations for the same diseases in the same patient were ignored and only the initial date of admission for a disease was used as a date of diagnosis for disease. In addition, we only counted 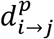 when the time between two admission dates was less than one year to filter out cases in which the diagnosis happened over a very long-term duration. A value of 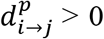 indicates the following: an initial admission for disease *i* occurred before the first admission for disease *j* in a patient *p* within one year. Then, the value of *δ_i→j_* was determined by the mean value of 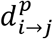 among the set of patients who were diagnosed with diseases *i* and *j* within one year. Thus, a value of *δ_i→j_* > 0 indicates that over half of admissions for disease *i* occurred before the admissions for disease *j* by one year among the patients who were diagnosed with both of these diseases. The statistical significance of disease pair co-occurrences (RA) and temporal directionality of diseases (*δ_i→j_*) were determined by using a binomial test (Benjamini–Hochberg FDR < 0.1) (Jensen et al., 2014). Finally, we used pairs of correlated diseases with time directionality whose mathematical relationships were statistically significant (RA > 1, FDR < 0.1; *δ_i→j_* ≠ 0, FDR < 0.1) for further analysis. We use the term ‘temporal disease correlation’ to describe this relationship.

### Disease trajectories (DAGs)

We joined multiple temporal disease correlations by concatenating temporal disease correlations into three or more steps of overall disease occurrences among patients (i.e., Disease 1 → 2 and Disease 2 → 3 to form Disease 1 → 2 → 3) (Jensen et al., 2014). Because our graph model only concedes a directional pair of temporal disease correlation, the result of concatenation is a directed acyclic graph (DAG), which means that the model presents a serial pattern of disease diagnoses by time without a loop (i.e., relapse of the same disease diagnoses again). A greedy approach was used to find subsequent steps in disease paths. Disease pairs were sorted in descending order according to patient counts. Pairs with overlapping diagnoses were found starting from the top of the list, and the number of patients following the full trajectory to death was counted. We stopped when a trajectory had no patients following it.

### Data validation and case review using USCF EHRs

We utilized EHRs from UCSF collected using the Epic system (Verona, WI) between 2011 and 2018, which includes inpatient and outpatient records for 836,461 unique individuals. The records were deidentified and contained no direct patient identifiers as defined in the Health Insurance Portability and Accountability Act (HIPAA). For the case review of schizophrenia and rhabdomyolysis patients, the analysis protocols were reviewed and approved by the UCSF institutional review board (approval number 17-22258).

## List of Supplementary Materials

Supplementary Movie S1

Supplementary Table S1

Supplementary Data S1

## Acknowledgements

We thank Jae Hyun Yoo for his work in developing the interactive data visualization of the trajectories and the showcase video. We thank Christina Mangurian, Keiichi Kodama, Dvir Aran, Kelly Zalocusky, and Seok Jong Yu for useful discussions.

## Author Contributions

H.P. and A.J.B. conceived the project and developed the algorithm of analysis pipeline. H.P. performed data processing and validation and H.P., M.J.K., N.R and U.M. performed data analysis. M.J.K. performed the patient case review. H.P. M.J.K and N.R. generated the figures and H.P./M.J.K. wrote the manuscript with critical input from D.H., M.S., B.C., U.M., S.B.C., and A.J.B.

## Data and Materials Availability

Inpatient records of over 10.2 million patients from California hospitals are available via material transfer agreement with the HCUP of the AHRQ (https://www.ahrq.gov/research/data/hcup/).

## References

American Academy of Family Physicians. JM, Marinides G, Wang GK. 1970. Rhabdomyolysis, American Family Physician. American Academy of Family Physicians.

Anttila V, Bulik-Sullivan B, Finucane HK, Walters RK, Bras J, Duncan L, Escott-Price V, Falcone GJ, Gormley P, Malik R, Patsopoulos NA, Ripke S, Wei Z, Yu D, Lee PH, Turley P, Grenier-Boley B, Chouraki V, Kamatani Y, Berr C, Letenneur L, Hannequin D, Amouyel P, Boland A, Deleuze J-F, Duron E, Vardarajan BN, Reitz C, Goate AM, Huentelman MJ, Kamboh MI, Larson EB, Rogaeva E, St George-Hyslop P, Hakonarson H, Kukull WA, Farrer LA, Barnes LL, Beach TG, Demirci FY, Head E, Hulette CM, Jicha GA, Kauwe JSK, Kaye JA, Leverenz JB, Levey AI, Lieberman AP, Pankratz VS, Poon WW, Quinn JF, Saykin AJ, Schneider LS, Smith AG, Sonnen JA, Stern RA, Van Deerlin VM, Van Eldik LJ, Harold D, Russo G, Rubinsztein DC, Bayer A, Tsolaki M, Proitsi P, Fox NC, Hampel H, Owen MJ, Mead S, Passmore P, Morgan K, Nöthen MM, Schott JM, Rossor M, Lupton MK, Hoffmann P, Kornhuber J, Lawlor B, McQuillin A, Al-Chalabi A, Bis JC, Ruiz A, Boada M, Seshadri S, Beiser A, Rice K, van der Lee SJ, De Jager PL, Geschwind DH, Riemenschneider M, Riedel-Heller S, Rotter JI, Ransmayr G, Hyman BT, Cruchaga C, Alegret M, Winsvold B, Palta P, Farh K-H, Cuenca-Leon E, Furlotte N, Kurth T, Ligthart L, Terwindt GM, Freilinger T, Ran C, Gordon SD, Borck G, Adams HHH, Lehtimäki T, Wedenoja J, Buring JE, Schürks M, Hrafnsdottir M, Hottenga J-J, Penninx B, Artto V, Kaunisto M, Vepsäläinen S, Martin NG, Montgomery GW, Kurki MI, Hämäläinen E, Huang H, Huang J, Sandor C, Webber C, Muller-Myhsok B, Schreiber S, Salomaa V, Loehrer E, Göbel H, Macaya A, Pozo-Rosich P, Hansen T, Werge T, Kaprio J, Metspalu A, Kubisch C, Ferrari MD, Belin AC, van den Maagdenberg AMJM, Zwart J-A, Boomsma D, Eriksson N, Olesen J, Chasman DI, Nyholt DR, Anney R, Avbersek A, Baum L, Berkovic S, Bradfield J, Buono R, Catarino CB, Cossette P, De Jonghe P, Depondt C, Dlugos D, Ferraro TN, French J, Hjalgrim H, Jamnadas-Khoda J, Kälviäinen R, Kunz WS, Lerche H, Leu C, Lindhout D, Lo W, Lowenstein D, McCormack M, Møller RS, Molloy A, Ng P-W, Oliver K, Privitera M, Radtke R, Ruppert A-K, Sander T, Schachter S, Schankin C, Scheffer I, Schoch S, Sisodiya SM, Smith P, Sperling M, Striano P, Surges R, Thomas GN, Visscher F, Whelan CD, Zara F, Heinzen EL, Marson A, Becker F, Stroink H, Zimprich F, Gasser T, Gibbs R, Heutink P, Martinez M, Morris HR, Sharma M, Ryten M, Mok KY, Pulit S, Bevan S, Holliday E, Attia J, Battey T, Boncoraglio G, Thijs V, Chen W-M, Mitchell B, Rothwell P, Sharma P, Sudlow C, Vicente A, Markus H, Kourkoulis C, Pera J, Raffeld M, Silliman S, Boraska Perica V, Thornton LM, Huckins LM, William Rayner N, Lewis CM, Gratacos M, Rybakowski F, Keski-Rahkonen A, Raevuori A, Hudson JI, Reichborn-Kjennerud T, Monteleone P, Karwautz A, Mannik K, Baker JH, O’Toole JK, Trace SE, Davis OSP, Helder SG, Ehrlich S, Herpertz-Dahlmann B, Danner UN, van Elburg AA, Clementi M, Forzan M, Docampo E, Lissowska J, Hauser J, Tortorella A, Maj M, Gonidakis F, Tziouvas K, Papezova H, Yilmaz Z, Wagner G, Cohen-Woods S, Herms S, Julià A, Rabionet R, Dick DM, Ripatti S, Andreassen OA, Espeseth T, Lundervold AJ, Steen VM, Pinto D, Scherer SW, Aschauer H, Schosser A, Alfredsson L, Padyukov L, Halmi KA, Mitchell J, Strober M, Bergen AW, Kaye W, Szatkiewicz JP, Cormand B, Ramos-Quiroga JA, Sánchez-Mora C, Ribasés M, Casas M, Hervas A, Arranz MJ, Haavik J, Zayats T, Johansson S, Williams N, Elia J, Dempfle A, Rothenberger A, Kuntsi J, Oades RD, Banaschewski T, Franke B, Buitelaar JK, Arias Vasquez A, Doyle AE, Reif A, Lesch K-P, Freitag C, Rivero O, Palmason H, Romanos M, Langley K, Rietschel M, Witt SH, Dalsgaard S, Børglum AD, Waldman I, Wilmot B, Molly N, Bau CHD, Crosbie J, Schachar R, Loo SK, McGough JJ, Grevet EH, Medland SE, Robinson E, Weiss LA, Bacchelli E, Bailey A, Bal V, Battaglia A, Betancur C, Bolton P, Cantor R, Celestino-Soper P, Dawson G, De Rubeis S, Duque F, Green A, Klauck SM, Leboyer M, Levitt P, Maestrini E, Mane S, De-Luca DM-, Parr J, Regan R, Reichenberg A, Sandin S, Vorstman J, Wassink T, Wijsman E, Cook E, Santangelo S, Delorme R, Rogé B, Magalhaes T, Arking D, Schulze TG, Thompson RC, Strohmaier J, Matthews K, Melle I, Morris D, Blackwood D, McIntosh A, Bergen SE, Schalling M, Jamain S, Maaser A, Fischer SB, Reinbold CS, Fullerton JM, Grigoroiu-Serbanescu M, Guzman-Parra J, Mayoral F, Schofield PR, Cichon S, Mühleisen TW, Degenhardt F, Schumacher J, Bauer M, Mitchell PB, Gershon ES, Rice J, Potash JB, Zandi PP, Craddock N, Ferrier IN, Alda M, Rouleau GA, Turecki G, Ophoff R, Pato C, Anjorin A, Stahl E, Leber M, Czerski PM, Edenberg HJ, Cruceanu C, Jones IR, Posthuma D, Andlauer TFM, Forstner AJ, Streit F, Baune BT, Air T, Sinnamon G, Wray NR, MacIntyre DJ, Porteous D, Homuth G, Rivera M, Grove J, Middeldorp CM, Hickie I, Pergadia M, Mehta D, Smit JH, Jansen R, de Geus E, Dunn E, Li QS, Nauck M, Schoevers RA, Beekman AT, Knowles JA, Viktorin A, Arnold P, Barr CL, Bedoya-Berrio G, Bienvenu OJ, Brentani H, Burton C, Camarena B, Cappi C, Cath D, Cavallini M, Cusi D, Darrow S, Denys D, Derks EM, Dietrich A, Fernandez T, Figee M, Freimer N, Gerber G, Grados M, Greenberg E, Hanna GL, Hartmann A, Hirschtritt ME, Hoekstra PJ, Huang A, Huyser C, Illmann C, Jenike M, Kuperman S, Leventhal B, Lochner C, Lyon GJ, Macciardi F, Madruga-Garrido M, Malaty IA, Maras A, McGrath L, Miguel EC, Mir P, Nestadt G, Nicolini H, Okun MS, Pakstis A, Paschou P, Piacentini J, Pittenger C, Plessen K, Ramensky V, Ramos EM, Reus V, Richter MA, Riddle MA, Robertson MM, Roessner V, Rosário M, Samuels JF, Sandor P, Stein DJ, Tsetsos F, Van Nieuwerburgh F, Weatherall S, Wendland JR, Wolanczyk T, Worbe Y, Zai G, Goes FS, McLaughlin N, Nestadt PS, Grabe H-J, Depienne C, Konkashbaev A, Lanzagorta N, Valencia-Duarte A, Bramon E, Buccola N, Cahn W, Cairns M, Chong SA, Cohen D, Crespo-Facorro B, Crowley J, Davidson M, DeLisi L, Dinan T, Donohoe G, Drapeau E, Duan J, Haan L, Hougaard D, Karachanak-Yankova S, Khrunin A, Klovins J, Kučinskas V, Lee Chee Keong J, Limborska S, Loughland C, Lönnqvist J, Maher B, Mattheisen M, McDonald C, Murphy KC, Murray R, Nenadic I, van Os J, Pantelis C, Pato M, Petryshen T, Quested D, Roussos P, Sanders AR, Schall U, Schwab SG, Sim K, So H-C, Stögmann E, Subramaniam M, Toncheva D, Waddington J, Walters J, Weiser M, Cheng W, Cloninger R, Curtis D, Gejman P V., Henskens F, Mattingsdal M, Oh S-Y, Scott R, Webb B, Breen G, Churchhouse C, Bulik CM, Daly M, Dichgans M, Faraone S V., Guerreiro R, Holmans P, Kendler KS, Koeleman B, Mathews CA, Price A, Scharf J, Sklar P, Williams J, Wood NW, Cotsapas C, Palotie A, Smoller JW, Sullivan P, Rosand J, Corvin A, Neale BM, Murray R. 2018. Analysis of shared heritability in common disorders of the brain. Science (80-) 360:eaap8757. doi:10.1126/science.aap8757

Ashley EA, Butte AJ, Wheeler MT, Chen R, Klein TE, Dewey FE, Dudley JT, Ormond KE, Pavlovic A, Morgan AA, Pushkarev D, Neff NF, Hudgins L, Gong L, Hodges LM, Berlin DS, Thorn CF, Sangkuhl K, Hebert JM, Woon M, Sagreiya H, Whaley R, Knowles JW, Chou MF, Thakuria J V, Rosenbaum AM, Zaranek AW, Church GM, Greely HT, Quake SR, Altman RB. 2010. Clinical assessment incorporating a personal genome. Lancet (London, England) 375:1525–35. doi:10.1016/S0140-6736(10)60452-7

Blair DR, Lyttle CS, Mortensen JM, Bearden CF, Jensen AB, Khiabanian H, Melamed R, Rabadan R, Bernstam E V, Brunak S, Jensen LJ, Nicolae D, Shah NH, Grossman RL, Cox NJ, White KP, Rzhetsky A. 2013. A nondegenerate code of deleterious variants in Mendelian loci contributes to complex disease risk. Cell 155:70–80. doi:10.1016/j.cell.2013.08.030

Camilo O, Goldstein LB. 2004. Seizures and epilepsy after ischemic stroke. Stroke 35:1769–75. doi:10.1161/01.STR.0000130989.17100.96

Chang K-Y, Wu Y-F. 2011. Aripiprazole-Associated Rhabdomyolysis in a Patient With Schizophrenia. J Neuropsychiatry Clin Neurosci 23:E51–E51. doi:10.1176/jnp.23.3.jnpe51

Consortium SWG of the PG, Ripke S, Neale BM, Corvin A, Walters JTR, Farh K-H, Holmans PA, Lee P, Bulik-Sullivan B, Collier DA, Huang H, Pers TH, Agartz I, Agerbo E, Albus M, Alexander M, Amin F, Bacanu SA, Begemann M, Jr RAB, Bene J, Bergen SE, Bevilacqua E, Bigdeli TB, Black DW, Bruggeman R, Buccola NG, Buckner RL, Byerley W, Cahn W, Cai G, Campion D, Cantor RM, Carr VJ, Carrera N, Catts S V., Chambert KD, Chan RCK, Chen RYL, Chen EYH, Cheng W, Cheung EFC, Chong SA, Cloninger CR, Cohen D, Cohen N, Cormican P, Craddock N, Crowley JJ, Curtis D, Davidson M, Davis KL, Degenhardt F, Favero J Del, Demontis D, Dikeos D, Dinan T, Djurovic S, Donohoe G, Drapeau E, Duan J, Dudbridge F, Durmishi N, Eichhammer P, Eriksson J, Escott-Price V, Essioux L, Fanous AH, Farrell MS, Frank J, Franke L, Freedman R, Freimer NB, Friedl M, Friedman JI, Fromer M, Genovese G, Georgieva L, Giegling I, Giusti-Rodríguez P, Godard S, Goldstein JI, Golimbet V, Gopal S, Gratten J, Haan L de, Hammer C, Hamshere ML, Hansen M, Hansen T, Haroutunian V, Hartmann AM, Henskens FA, Herms S, Hirschhorn JN, Hoffmann P, Hofman A, Hollegaard M V., Hougaard DM, Ikeda M, Joa I, Julià A, Kahn RS, Kalaydjieva L, Karachanak-Yankova S, Karjalainen J, Kavanagh D, Keller MC, Kennedy JL, Khrunin A, Kim Y, Klovins J, Knowles JA, Konte B, Kucinskas V, Kucinskiene ZA, Kuzelova-Ptackova H, Kähler AK, Laurent C, Keong JLC, Lee SH, Legge SE, Lerer B, Li M, Li T, Liang K-Y, Lieberman J, Limborska S, Loughland CM, Lubinski J, Lönnqvist J, Jr MM, Magnusson PKE, Maher BS, Maier W, Mallet J, Marsal S, Mattheisen M, Mattingsdal M, McCarley RW, McDonald C, McIntosh AM, Meier S, Meijer CJ, Melegh B, Melle I, Mesholam-Gately RI, Metspalu A, Michie PT, Milani L, Milanova V, Mokrab Y, Morris DW, Mors O, Murphy KC, Murray RM, Myin-Germeys I, Müller-Myhsok B, Nelis M, Nenadic I, Nertney DA, Nestadt G, Nicodemus KK, Nikitina-Zake L, Nisenbaum L, Nordin A, O’Callaghan E, O’Dushlaine C, O’Neill FA, Oh S-Y, Olincy A, Olsen L, Os J Van, Pantelis C, Papadimitriou GN, Papiol S, Parkhomenko E, Pato MT, Paunio T, Pejovic-Milovancevic M, Perkins DO, Pietiläinen O, Pimm J, Pocklington AJ, Powell J, Price A, Pulver AE, Purcell SM, Quested D, Rasmussen HB, Reichenberg A, Reimers MA, Richards AL, Roffman JL, Roussos P, Ruderfer DM, Salomaa V, Sanders AR, Schall U, Schubert CR, Schulze TG, Schwab SG, Scolnick EM, Scott RJ, Seidman LJ, Shi J, Sigurdsson E, Silagadze T, Silverman JM, Sim K, Slominsky P, Smoller JW, So H-C, Spencer CA, Stahl EA, Stefansson H, Steinberg S, Stogmann E, Straub RE, Strengman E, Strohmaier J, Stroup TS, Subramaniam M, Suvisaari J, Svrakic DM, Szatkiewicz JP, Söderman E, Thirumalai S, Toncheva D, Tosato S, Veijola J, Waddington J, Walsh D, Wang D, Wang Q, Webb BT, Weiser M, Wildenauer DB, Williams NM, Williams S, Witt SH, Wolen AR, Wong EHM, Wormley BK, Xi HS, Zai CC, Zheng X, Zimprich F, Wray NR, Stefansson K, Visscher PM, Consortium WTC-C, Adolfsson R, Andreassen OA, Blackwood DHR, Bramon E, Buxbaum JD, Børglum AD, Cichon S, Darvasi A, Domenici E, Ehrenreich H, Esko T, Gejman P V., Gill M, Gurling H, Hultman CM, Iwata N, Jablensky A V., Jönsson EG, Kendler KS, Kirov G, Knight J, Lencz T, Levinson DF, Li QS, Liu J, Malhotra AK, McCarroll SA, McQuillin A, Moran JL, Mortensen PB, Mowry BJ, Nöthen MM, Ophoff RA, Owen MJ, Palotie A, Pato CN, Petryshen TL, Posthuma D, Rietschel M, Riley BP, Rujescu D, Sham PC, Sklar P, Clair DS, Weinberger DR, Wendland JR, Werge T, Daly MJ, Sullivan PF, O’Donovan MC. 2014. Biological insights from 108 schizophrenia-associated genetic loci. Nature 511:421–427. doi:10.1038/nature13595

Finkelstein J, Cha E, Scharf SM. 2009. Chronic obstructive pulmonary disease as an independent risk factor for cardiovascular morbidity. Int J Chron Obstruct Pulmon Dis 4:337–49.

Goldman LS. 1999. Medical illness in patients with schizophrenia. J Clin Psychiatry 60 Suppl 21:10–5.

Greenaway M, Elbe D. 2009. Focus on aripiprazole: a review of its use in child and adolescent psychiatry. J Can Acad Child Adolesc Psychiatry 18:250–60.

Hidalgo CA, Blumm N, Barabasi AL, Christakis NA. 2009. A dynamic network approach for the study of human phenotypes. PLoS Comput Biol 5:e1000353. doi:10.1371/journal.pcbi.1000353

Jensen AB, Moseley PL, Oprea TI, Ellesøe SG, Eriksson R, Schmock H, Jensen PB, Jensen LJ, Brunak S. 2014. Temporal disease trajectories condensed from population-wide registry data covering 6.2 million patients. Nat Commun 5:4022. doi:10.1038/ncomms5022

Life I of M (US) C on C at the E of, Field MJ, Cassel CK. 1997. A Profile of Death and Dying in America. National Academies Press (US).

Martin GS. 2012. Sepsis, severe sepsis and septic shock: changes in incidence, pathogens and outcomes. Expert Rev Anti Infect Ther 10:701–6. doi:10.1586/eri.12.50

Melli G, Chaudhry V, Cornblath DR. 2005. Rhabdomyolysis: an evaluation of 475 hospitalized patients. Medicine (Baltimore) 84:377–85.

Murdoch TB, Detsky AS, S G, JL B, D C, C S, HJ M, BY R, PB J. 2013. The Inevitable Application of Big Data to Health Care. JAMA 309:1351. doi:10.1001/jama.2013.393

O’Malley KJ, Cook KF, Price MD, Wildes KR, Hurdle JF, Ashton CM. 2005. Measuring diagnoses: ICD code accuracy. Health Serv Res 40:1620–39. doi:10.1111/j.1475-6773.2005.00444.x

Park J, Lee DS, Christakis NA, Barabasi AL. 2009. The impact of cellular networks on disease comorbidity. Mol Syst Biol 5:262. doi:msb200916 [pii]10.1038/msb.2009.16

Prasad S, Bhatia T, Kukshal P, Nimgaonkar VL, Deshpande SN, Thelma BK. 2017. Attempts to replicate genetic associations with schizophrenia in a cohort from north India. npj Schizophr 3:28. doi:10.1038/s41537-017-0030-8

Prior MJ, Foletta VC, Jowett JB, Segal DH, Carless MA, Curran JE, Dyer TD, Moses EK, McAinch AJ, Konstantopoulos N, Bozaoglu K, Collier GR, Cameron-Smith D, Blangero J, Walder KR. 2010. The characterization of Abelson helper integration site–1 in skeletal muscle and its links to the metabolic syndrome. Metabolism 59:1057–1064. doi:10.1016/j.metabol.2009.11.002

Steiner C, Elixhauser A, Schnaier J. The healthcare cost and utilization project: an overview. Eff Clin Pract 5:143–51.

Yeo Y, Gwack J, Kang S, Koo B, Jung SJ, Dhamala P, Ko K-P, Lim Y-K, Yoo K-Y. 2013. Viral hepatitis and liver cancer in Korea: an epidemiological perspective. Asian Pac J Cancer Prev 14:6227–31.

